# Access COI barcode efficiently using high throughput Single-End 400 bp sequencing

**DOI:** 10.1101/498618

**Authors:** Chentao Yang, Shangjin Tan, Guangliang Meng, David G. Bourne, Paul A. O’Brien, Junqiang Xu, Sha Liao, Ao Chen, Xiaowei Chen, Shanlin Liu

## Abstract

1. Over the last decade, the rapid development of high-throughput sequencing platforms has accelerated species description and assisted morphological classification through DNA barcoding. However, constraints in barcoding costs led to unbalanced efforts which prevented accurate taxonomic identification for biodiversity studies.
2. We present a high throughput sequencing approach based on the HIFI-SE pipeline which takes advantage of Single-End 400 bp (SE400) sequencing data generated by BGISEQ-500 to produce full-length Cytochrome *c* oxidase subunit I (COI) barcodes from pooled polymerase chain reaction amplicons. HIFI-SE was written in Python and included four function modules of *filter, assign, assembly* and *taxonomy*.
3. We applied the HIFI-SE to a test plate which contained 96 samples (30 corals, 64 insects and 2 blank controls) and delivered a total of 86 fully assembled HIFI *COI* barcodes. By comparing to their corresponding Sanger sequences (72 sequences available), it showed that most of the samples (98.61%, 71/72) were correctly and accurately assembled, including 46 samples that had a similarity of 100% and 25 of ca. 99%.
4. Our approach can produce standard full-length barcodes cost efficiently, allowing DNA barcoding for global biomes which will advance DNA-based species identification for various ecosystems and improve quarantine biosecurity efforts.

## Introduction

Since it was first proposed by Hebert *et al. (Hebert, Cywinska & Ball 2003)*, DNA barcoding has attracted global synergistic efforts resulting in well-curated and centralized reference databases. The Barcode of Life Data systems (BOLD) (Ratnasingham & Hebert 2007), for example, has been growing into a repository of greater than 5.8 million barcodes representing 282,738 species (accessed in Nov. 2018). The strength of DNA barcoding includes its application across all stages of life, from lava to adult and even predator feces (Symondson 2002; Valentini, Pompanon & Taberlet 2009) and stomach contents (Krehenwinkel *et al*. 2017). This, along with the ease of barcoding accessibility and analysis, has led to its use in a wide spectrum of scientific and commercial areas, such as cryptic species discovery (Bączkiewicz *et al*. 2017), biodiversity monitoring (Bohmann *et al*. 2014; Tang *et al*. 2015; Thomsen & Willerslev 2015), conservation biology (Krishnamurthy, Francis & conservation 2012), inspection of illegal trade of endangered species (Collins *et al*. 2012) and discovery of illegal ingredients in medicine (Coghlan *et al*. 2012).

Even though barcode sequences have been accumulating rapidly in the last decade, the available reference databases are limited by poor and biased spatial coverage and skewed taxonomic coverage (Yoccoz 2012). Biodiversity initiatives typically suffer from insufficient funding for DNA-based taxonomy work, and scientists have been attempting to achieve barcode sequences in a cost-efficient way via high-throughput sequencing (HTS) platforms. However, these methods, owing to their read lengths, only deliver a fraction of the standard barcode (Meier *et al*. 2016), or require either extra laboratory workloads of multiple rounds of PCRs (Shokralla *et al*. 2015; Cruaud *et al*. 2017), or an extra K-mer based assembly step leading to accuracy uncertainty (Liu *et al*. 2017). Long reads from the Single Molecular Real Time (SMRT) sequencing platform or nanopore platform may achieve reliable standard barcode sequences, however, at a higher cost (Liu *et al*. 2017; Hebert *et al*. 2018). Standard barcode (*COI*, cytochrome *c* oxidase I gene) for animals with its flanking primers and tags is ca. 700 bp in length, the HTS platform offers significant advantages since it allows for accurate *COI* barcode assembly by connecting the 5′ and 3′ reads provided the HTS platform can generate reads of a length ≥ 400 bp, with a minimum overlap of ~ 80 bp.

The BGISEQ-500 platform has launched a new test sequencing kit capable of single-end 400 bp sequencing (SE400), which offers a simple and reliable way to achieve DNA barcodes efficiently. In this study, we explore the potential of the BGISEQ-500 SE400 sequencing in DNA barcode reference construction and provide an updated HIFI-SE barcode software package that can generate *COI* barcode assemblies using HTS reads of a length of 400 bp.

## Overview of the HIFI-SE barcode pipeline

### Experimental process

#### 1) DNA preparation

DNA of each well in the plate should be extracted separately before PCR. The Glass Fiber Plate method (Ivanova, Dewaard & Hebert 2006) is recommended because of relatively high efficiency and low cost.

#### 2) PCR amplification

96 paired tags were added to both ends of the common *COI* barcode primer set (LCO1490 and HCO2198 (O. Folmer 1994)) (Supplementary Table S1). Each tag was 5 bp in length and had ≥ 2 bp difference from each other. Each PCR reaction (25 μL) contained 1 μL DNA template, 16.2 μL molecular biology grade water, 2.5 μL 10× buffer (Mg^2+^ plus), 2.5 μL dNTP mix (10 mM), 1 μL each forward and reverse primers (10 mM), and 0.3 μL TaKaRa Ex Taq polymerase (5 U/μL) (Takara, Dalian, China). The amplification program included a thermocycling profile of 94°C for 60s, 5 cycles of 94°C for 30 s, 45 °C for 40 s, and an extension at 72 °C for 60 s, followed by 35 cycles of 94 °C for 30 s, 51 °C for 40 s, and 72 °C for 60 s, with a final extension at 72 °C for 10 min, and a final on-hold at 12 °C.

#### 3) Library construction and sequencing

One microliter of each amplicon was mixed together and subsequently sent to BGI-Shenzhen for library preparation and sequencing (BGISEQ-500 SE400 module) following the BGISeq-500 library construction protocol (Supplementary File 1).

### Bioinformatics

The software package is written in Python and is deposited in PyPI (https://pypi.org/project/HIFI-SE/), consisting of three main function modules of *‘filter’, ‘assign’*, and *‘assembly’* (Fig. 2). Full functions and a tutorial are detailed in the software manual (Supplementary File 2)

**Fig. 1:**
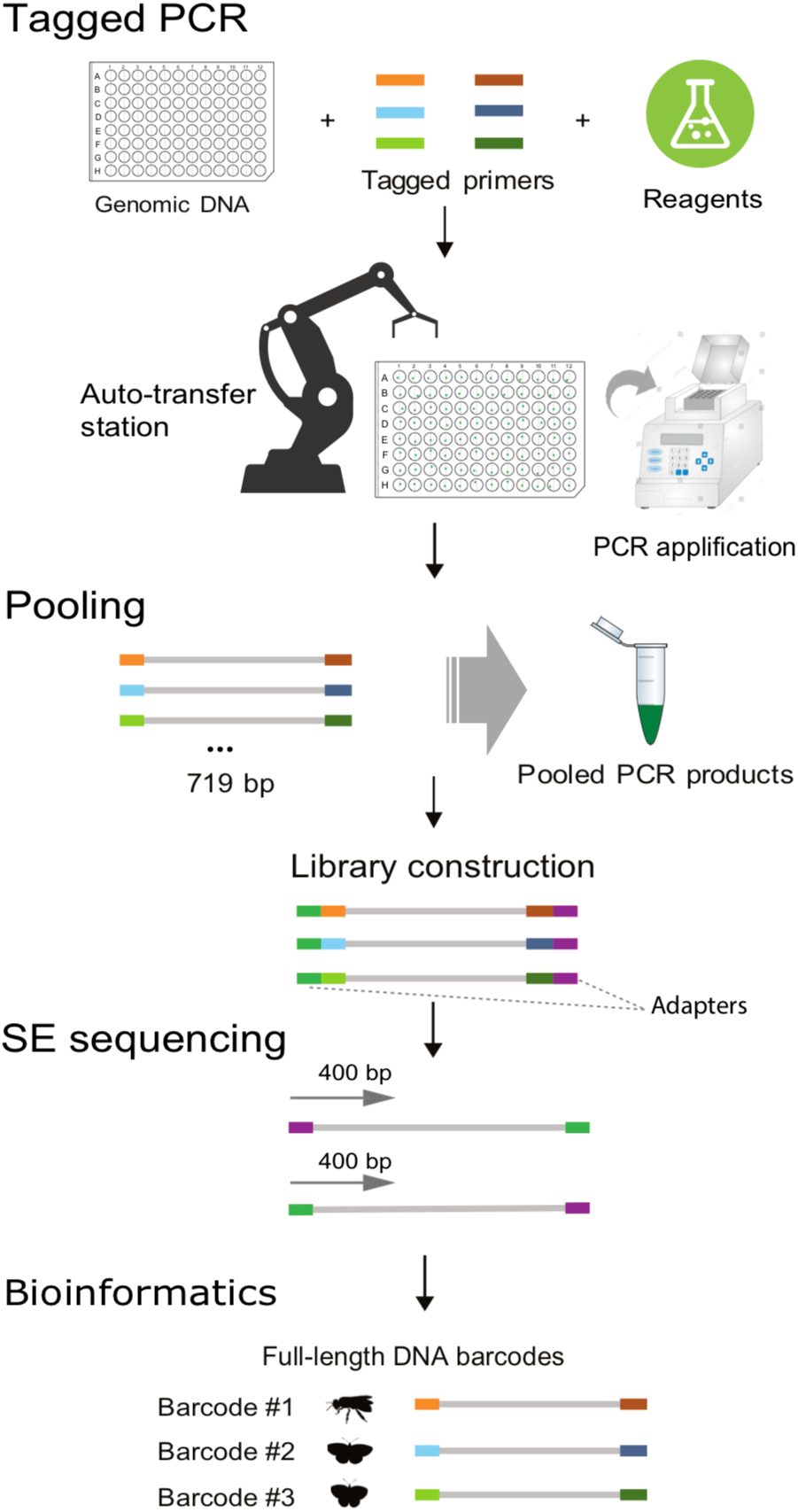
Schematic illustration of the HIFI-SE barcode pipeline.

**Fig. 2:**
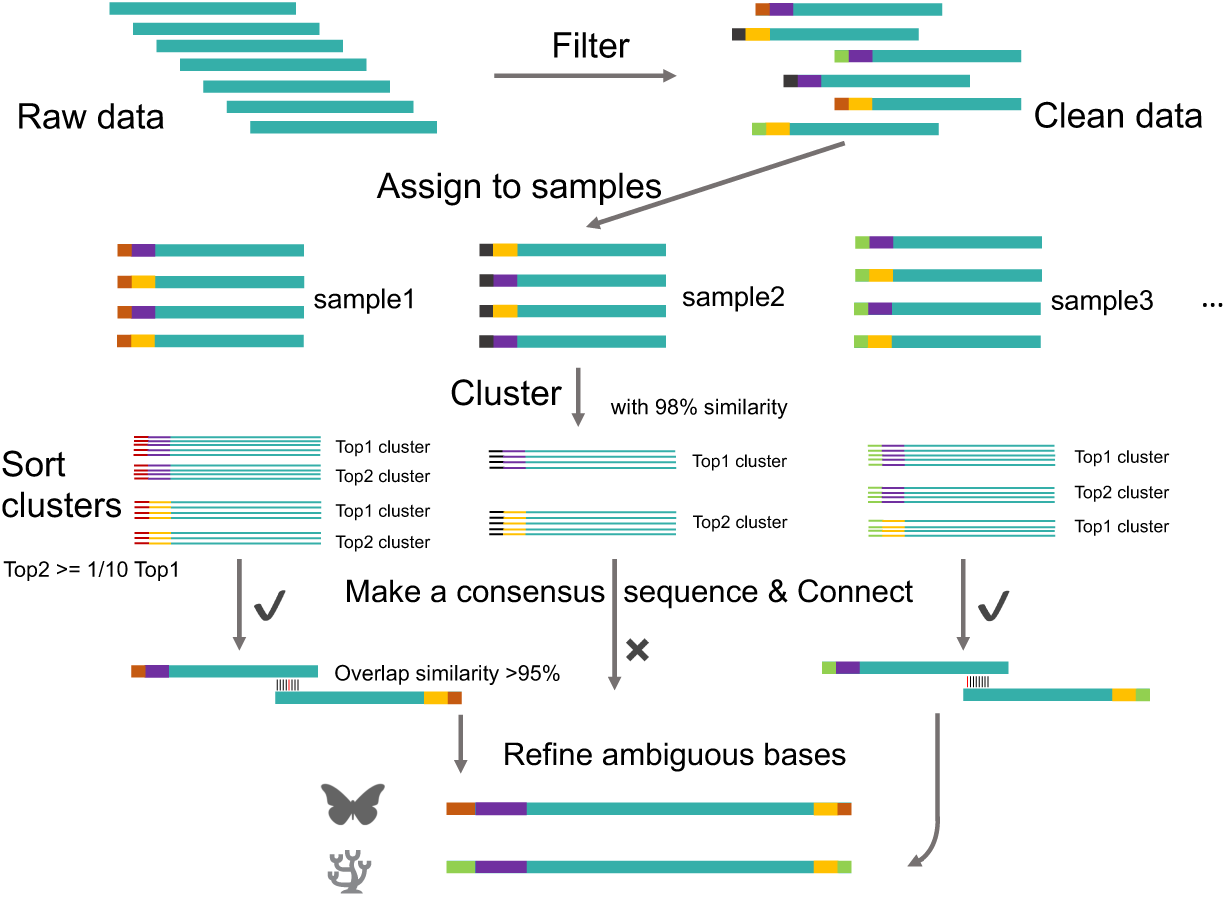
HIFI-SE barcode assembly pipeline. The colored bars from left to right represent tags, primers (purple for 5′ end and orange for 3′ end) and barcode sequences, respectively.

#### 1) Data filtering

It removed low quality reads of 1) reads with ambiguous bases; 2) reads with an expected error number *E\** >10 with *E\** being calculated using a formula of 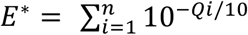, where n represents sequence length and *Qi* represents base quality of the *i^th^* base on reads.

#### 2) Read assignment

Reads were demultiplexed by index and then classed to the 5′ and 3′ ends by primer sequences. Both require a 100% identity.

#### 3) Full-length *COI* barcode assembly

For each end, sequences were first clustered at a 98% similarity using VSEARCH (v2.8.0) (Rognes *et al*. 2016). Subsequently, a consensus sequence was built from the most abundant cluster (cluster 1) supported by ≥ 5 sequences. Sequences from the second most abundant cluster were also retained to identify potential symbionts or parasites if containing sequence numbers > 1/10 of that in cluster 1.

Full-length *COI* barcodes were assembled by connecting the consensus sequences of the 5′ and 3′ ends with an overlap ≥ 80 bp and similarity ≥ 95%. Discrepancies were determined based on the base frequency in sequences from both ends. The assemblies with correct amino acid translation and a length of > 650 bp were output as final results. If an assembly failed with the default options, users can run another round with an additional parameter – checking for amino acid translation before clustering (supplemental File 2).

#### 4) Taxonomy identification in BOLD

The HIFI-SE pipeline provides an optional step (*Taxonomy*) to verify the taxonomic information of assembled sequences. It can automatically submit assemblies to the BOLD system and grab the taxonomic information from the returned searches. Currently, it supports searching of the animal, fungi and plant databases and outputs a user-defined number of BOLD items for each sequence.

## Example analysis

### Materials and methods

Specimens used in this study contained 30 corals, 64 insects and 2 blank controls (Supplementary Table S2). Corals were sampled from the Great Barrier Reef using hammer and chisel and kept in running seawater until processing. Insect samples were randomly chosen from collections from the Laohegou Natural Reserve, Sichuan Province, China. Coral tissue was removed from the skeleton using pressurized air from a blow gun into a ziplock bag containing 10ml of calcium magnesium free artificial seawater (CMFASW; NaCl 26.2 g, KCl 1 g, NaHCO_3_, Milli-Q H_2_O 1 L). Approximately 0.05 g of coral tissue pellet was used for DNA extraction using the PowerBiofilm DNA Isolation Kit (QIAGEN Pty Ltd, Australia) following the manufacturers protocol. Insect genomic DNA was extracted using the Glass Fiber Plate method (Ivanova, Dewaard & Hebert 2006) following the manufacturer’s protocol.

Before pooling for HTS sequencing, all the PCR products were sent to BGI-Shenzhen for Sanger sequencing from both 5′ and 3′ ends on an ABI 3730XL platform. A total of 73 sequences were successfully assembled using Geneious (Kearse *et al*. 2012) and served as a reference database to evaluate the accuracy and efficiency of the HIFI-SE pipeline. The 21 failed samples (excluding 2 blanks) were referred to as “Barcode failed” samples, and those failures can be attributed to the excessive non-targeted short PCR co-amplifications of a length of ca. 400 bp (Fig. S1 and detailed in the following Discussion part).

To evaluate the accuracy of HIFI-SE, barcodes obtained via HIFI-SE were aligned to their Sanger references using MUSCLE (Edgar 2004) and then checked for similarities between each. We subsequently aligned the demultiplexed reads to their corresponding HIFI-SE assemblies using BWA (Version: 0.7.17-r1188) (Li & Durbin 2009) to examine read support for sites at which the HIFI-SE and Sanger generated barcodes were different.

## Results

A total of 12,745,067 SE400 reads were retained after quality control. Around 77.9% (9,870,823) of reads were assigned to their corresponding samples as either 5′ or 3′ end. The number of sequences of each sample varied markedly, ranging from 303 to 585,609, with Sanger “barcode failed” samples possessing lower but insignificant number of reads (Fig. S1). A total of 86 barcode sequences including 63 insect samples and 23 coral samples were achieved using our pipeline with 14 out of the 21 Sanger “barcode failed” samples being successfully recovered, leading to an overall success rate of 91.5% (Fig. 3). There was also one sample that had a Sanger reference missed in the HIFI-SE assemblies.

**Fig. 3:**
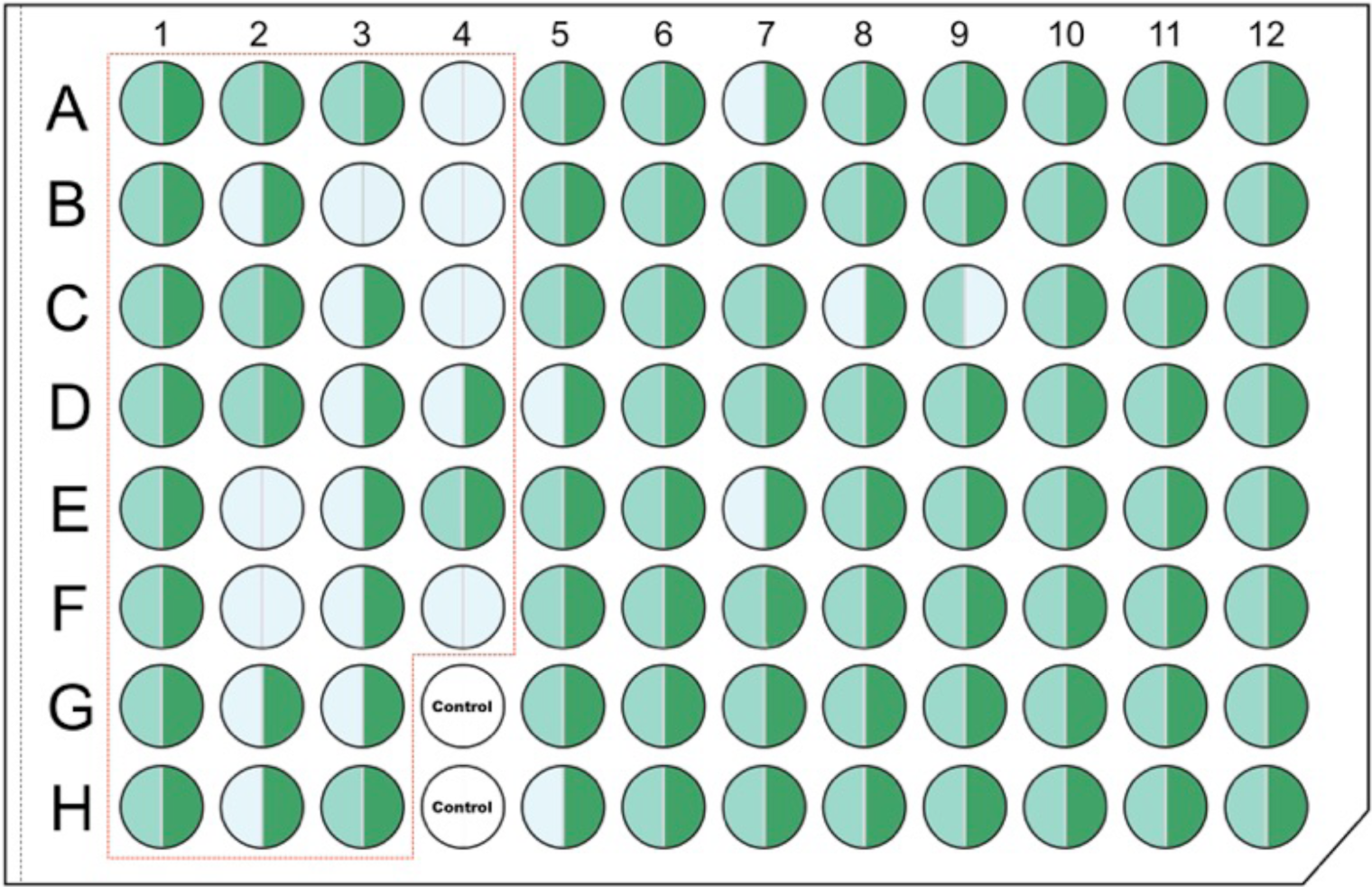
Results of Sanger sequencing (left semicircle) and HIFI-SE barcode assemblies (right semicircle) arranged in a 96-well plate. Gray represents failure; light and dark green represent success of Sanger and HIFI-SE respectively. Coral samples are arranged in wells from A01 to F04 (framed by the red tetragon). Insects are arranged in wells from A05 to H12.

Compared to the Sanger reference sequences (72 sequences available), HIFI-SE assemblies showed high-score matches for vast majority of the samples (98.61%, 71/72), including 46 samples that had a similarity of 100% and 25 of around 99% (Supplementary Table S3). Only one sample that showed a high dissimilarity score to its Sanger reference was demonstrated to be cross contamination from samples on the same plate. Read alignment results showed that the sites on HIFI-SE assemblies at which mismatches occurred were supported by high read coverage, confirming the accurate recovery of HIFI-SE (Fig. S2). In addition, HIFI-SE also identified a total of 40 ambiguous sites in the Sanger reference to specific nucleotides and revealed the heteroplasmy states in some samples (Fig. S2).

## Discussion

Despite the importance of biodiversity in ecosystem functioning (Tilman, Isbell & Cowles 2014), global biodiversity continues to be lost at an unprecedented rate due to climate change and human activities (Kerr & Currie 1995). DNA barcoding (Hebert, Cywinska & Ball 2003), has proven effective in accelerating the collection of biodiversity inventories over large geographic and temporal scales which benefits both researchers and also policy-makers focused on maintaining functioning ecosystems (Molnar *et al*. 2008). However, it is reported that hundreds of thousands of dollars are required to establish a DNA barcode reference database for a regular ecology study (Cameron, Rubinoff & Will 2006), and ca. $1 billion for sequencing to complete global barcode registration (Marshall 2005). The burgeoning massive parallel sequencing techniques drive the cost per nucleotide base down dramatically (Von Bubnoff 2008) and inspired multifarious approaches to obtain barcode sequences via HTS platforms (Liu *et al*. 2013; Liu *et al*. 2017; Hebert *et al*. 2018). The HIFI-SE pipeline which takes advantage of HTS reads as long as 400 bp, provides an easy, simple and cost effective (an average cost of $1 USD per barcode) approach to generate barcode sequences from a large number of samples. The 400 bp reads enable an overlap length of ca. 80 bp for most animal *COI* barcode sequences by sequencing both 5′ and 3′ ends, and the plain data process step – assembly by overlapping, can simplify the barcode assembly process by circumventing the *de Brujin* graph algorithm which is time-consuming and computationally intensive (Li *et al*. 2012) and can hardly avoid erroneous pathing when deal with intricate scenarios.

Two taxonomic groups, corals and insects, were included to demonstrate the effectiveness of this approach. The results showed that insects delivered higher sample recovery ratios (63 out of 64 samples) compared to corals (23 out of 30 samples). The relatively lower efficiency of coral can be attributed to the biased performance of primer set LCO1490 and HCO2198 in corals (Geller *et al*. 2013). It shows the necessity to promote primer design to fit more various phylogenetic lineages in spite of the high sensitivity of HTS methods. The primer’s inadequacy for coral was also reflected by excessive short co-amplicons (400~500bp) detected in 16 out of 21 Sanger “Barcode failed” samples (Fig. S1), which might be derived from Nuclear Mitochondrial DNA Segment (Numt) and in turn affect the recovery success of their barcode sequences via both the Sanger sequencing and HIFI-SE pipeline. Besides, coral is well known for being difficult in DNA extraction and tends to degrade quite rapidly for lots of species (Neigel, Domingo & Stake 2007). Thus, their DNA quality may also contribute to the short co-amplicons. It also reveals the strength of our approach in dealing with those samples that are difficult to work with using the traditional method. In addition, we also noticed one assembly (E08 in Supplementary Table S3) that showed low similarity to its corresponding Sanger reference was actually cross contamination from another cell (C11 or H12 in Fig. 3). Since we mixed PCR reagents and PCR products using an auto transfer station (Hamilton Microlab^®^ STAR) and sample E08 only contained a read number of 1,000, we believe this contamination event could result from pipette failure on the auto transfer station during sample transfer (occasionally happens), and a subsequent tag hopping from other samples during library construction and sequencing.

In summary, the HIFI-SE pipeline requires straightforward processing in both sequencing preparation and data analysis, and holds great potential to on one hand further reduce per unit cost of DNA barcoding while on the other increase the efficiency and accuracy of obtained barcodes. This is achieved by increasing the throughput capacity via increasing tag length to allow more index combinations, and pooling amplicons using different primer sets. In addition, although we used the *COI* barcode for demonstration, our pipeline is expected to fit other marker genes with a length of 600-750bp well (e.g. V1-V4, V3-V6, and V5-V9 of 16S rRNA gene). Therefore, this new approach can produce standard full-length barcodes cost efficiently, allowing initiatives targeted at DNA barcoding biomes more foreseeable and thereby improving our understanding of the biodiversity of our global ecosystems or improving DNA based biosecurity programs.

## Supporting information

## Author contributions

C.Y. and SL.L. conceived the idea and designed the methodology; C.Y. and G.M. developed the program; D.G.B and P.A.O. collected the coral samples and extracted DNA. C.Y. and S.T. collected, analyzed the data and drafted the manuscript; J.X., SH.L., A.C. and X.C. conducted the library construction and sequencing. SL.L. revised the manuscript and all authors approved for final publication.

## Acknowledgements

We thank Dr. Ding Yang from China Agricultural University for contributing samples. We would like to thank Guojie Zhang and Qiye Li for sample and SE400 sequencing coordination. This research was supported by Shenzhen Municipal Government of China (NO. JCYJ20170817150755701) and Shenzhen Peacock Plan (No. KQTD20150330171505310). Authors have no conflict of interest to declare.

## Data accessibility

The data reported in this study are available in the CNGB Nucleotide Sequence Archive (CNSA: https://db.cngb.org/cnsa; accession number CNP0000195) and the EMBL repository (PRJEB29212, ERP111495).

## Supporting Information

Supplementary Table S1. Sequence of the tagged primers

Supplementary Table S2. Sample Information

Supplementary Table S3. Accuracy results of HIFI-SE barcodes compared with Sanger

Supplementary Data 1. Results of HIFI-SE400 Barcodes

Supplementary Data 2. Results of Sanger barcodes.

Supplementary File 1. Library construction protocol of BGISEQ-500 SE400 module.

Supplementary File 2. The manual of HIFI-SE package.

**Fig. S1.**
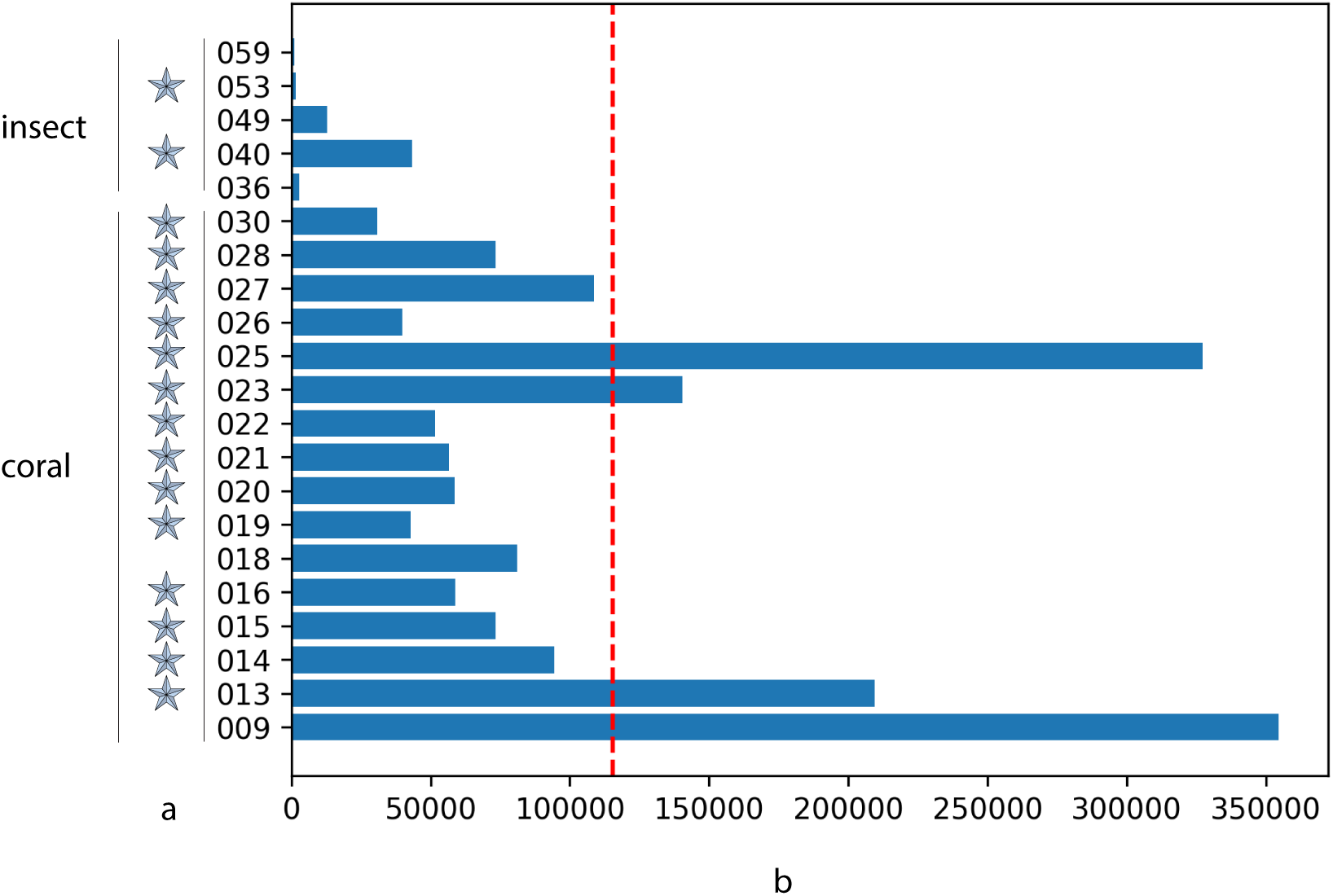
Read counts of the Sanger barcode failed samples. Stars indicate samples of which short amplicon(s) was detected in the HIFI assemblies. Short amplicons are those clusters of abundance >10 and of length < 600bp. The bar plot demonstrates the number of assigned reads for the barcode failed samples. The red dashed line shows the average value of all the successful samples and no significant difference was detected between the two groups (P value of 0.232, Student’s t-Test)

**Fig. S2.**
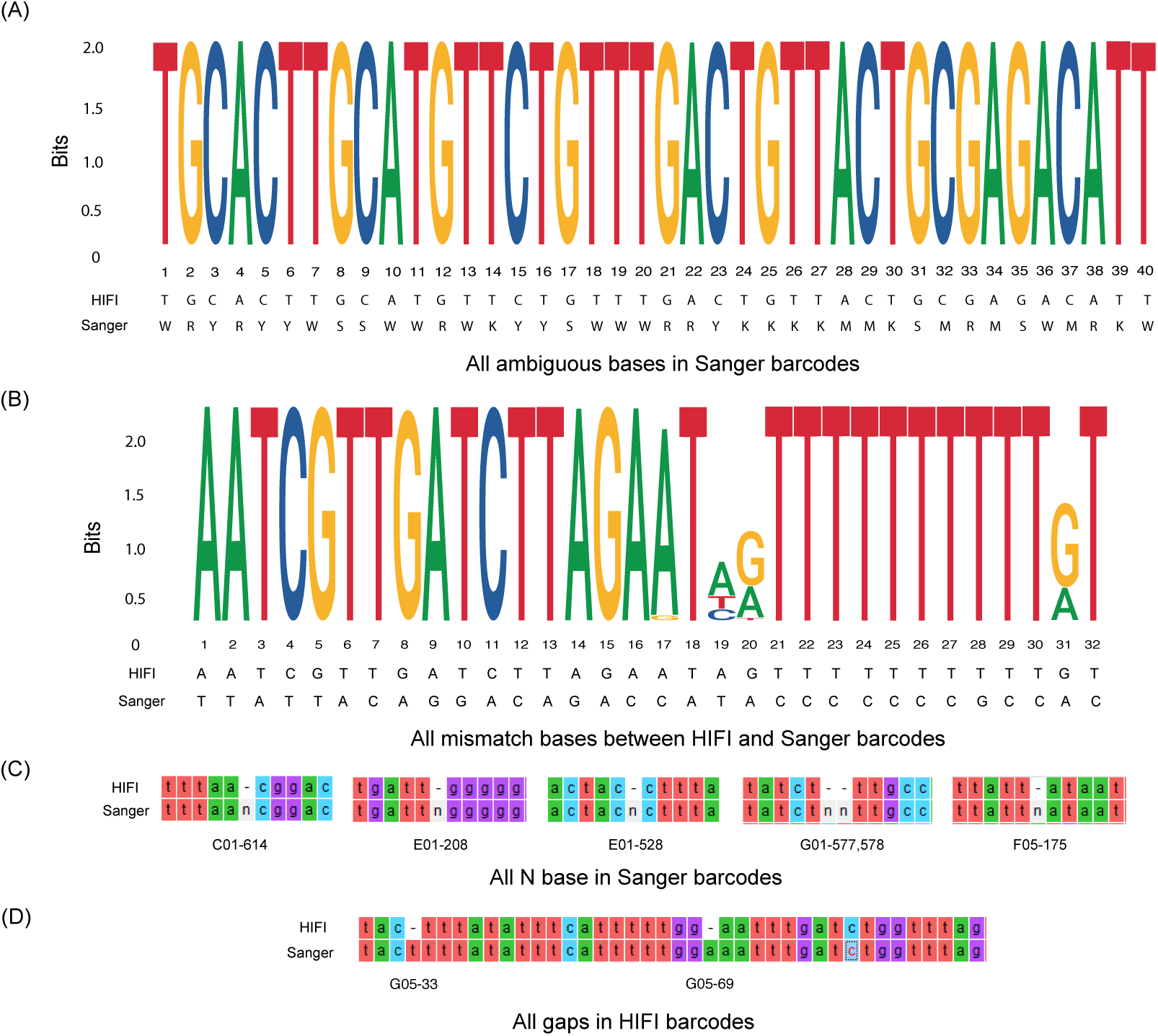
Discrepancies between Sanger sequences and HIFI-SE barcodes. Entropy weight was calculated based on the strength of read depth by aligning the SE400 reads onto the assembled HIFI-SE barcodes, showing differences between ambiguous Sanger base-calling and specific nucleotide identified in HIFI-SE barcodes (A) and potential heteroplasmy (B). In additional, several N bases were present of insertion in Sanger sequence (C), also two N bases in HIFI sequences (D).

