## Supplementary material for "Access COI barcode efficiently using high throughput Single-End 400 bp sequencing": Supplementary File 1.pdf

### DNA preparation

1) Corals

Coral tissue was removed from the skeleton using pressurized air from a blow gun into a ziplock bag containing 10ml of calcium magnesium free artificial seawater (CMFASW; NaCl 26.2g, KCl 1g, $NaHCO_3$, Milli-Q $H_2O$ 1L). Coral tissue blastate was aliquoted into 2ml microfuge tubes and pelleted in a fixed angle centrifuge at 10,000g for 10 min. Pellets were either snap frozen and stored at -80°C or preserved in 1ml salt saturated dimethyl sulfoxide (DMSO buffer; 23.265g disodium EDTA, 50ml 1M NaOH, 150ml Milli-Q $H_2O$, 50ml DMSO, NaCl till saturation) and stored at -20°C. Approximately 0.05g of coral tissue pellet was then used for DNA extraction using the PowerBiofilm DNA Isolation Kit (QIAGEN Pty Ltd, Australia) following the manufacturers protocol.

2) Insects

Individual Genomic DNA could be extracted using the Glass Fiber Plate method following manufacturer's protocol or other existing method (Ivanova, Dewaard & Hebert 2006).

### Library construction

This beta version protocol is for BGISEQ-500 library construction so as to adapt Single-End 400 bp sequencing module. Compared to the standard library construction protocol of BGISEQ-500 WGS module, this protocol removed DNA fragmentation and fragment selection steps.

### 1) DNA input and Homogenization

No need to fragment DNA (PCR production is around 700bp length).

1) Use double-strand DNA quantification kit such as Qubit® dsDNA HS Assay Kit or Quant-iT™ PicoGreen® dsDNA Assay Kit and quantify the sample as per the instructions of the quantification kit.
2) Remove 50 ng of sample (calculated based on its concentration) to a new 0.2 mL PCR tube, then add NF water to final volume of 40 μL.

### 2. End Repair and Tailing

1) Prepare the mixture as follows in PCR tube (do not vortex enzymes):

| Components | Volume |
|---|---|
| DNA | 40μL |
| ERAT Buffer | 7.1μL |
| ERAT Enzyme | 2.9μL |
| Total | 50μL |

2) Mix well by gently pipetting (Do not mix by vortexing), concentrate the reaction liquid to tube bottom by brief centrifugation.
3) Place the PCR tube containing the reaction mixture of above step in a Thermal Cycler, and initiate the reaction as per the following conditions:

| Temperature | Time |
|---|---|
| Heated lid | On |
| 37℃ | 30 min |
| 65℃ | 15 min |
| 4℃ | Hold |

### 3. Ligate Adapters

1) Add 5 μL of Adapter Mix to above PCR tube and mix well by pipetting. Now 16 Adapter Mix are available, 8 libraries in one lane strategy, every sample with 4 different barcodes.
2) Prepare the following reaction mixture (Note: Ligation Buffer is viscous, pipette slowly):

| Components | Volume |
|---|---|
| Ligation Buffer | 23.4 μL |
| Ligation Enzyme | 1.6 μL |
| Total | 25 μL |

3) Add 25 μL of above reaction mixture to the reaction solution containing adapters from above step.
4) Place the tube in a Thermal Cycler, then initiate reaction as per following condition:

| Temperature | Time |
|---|---|
| Heated lid | On |
| 23℃ | 30 min |
| 4℃ | Hold |

5) After ligation, add 20 μL TE to final volume of 100 μL, then transfer entire volume to a non-stick tube containing 50 uL of room temperature AMPure beads and mix by slow pipetting 10 times to avoid bubble formation.

### 4. Purify Ligated DNA

1) Incubate at room temperature for 5 min.
2) After brief centrifugation, place the non-stick tube on the magnet for 2 min until the liquid clears, remove and discard the supernatant with a pipette:
3) Add 500 μL of fresh 80% ethanol, while the tube remains on the magnet, then, rotate the tubes in the rack by half turns 4 times to wash the beads. Carefully remove and discard the supernatant.
4) Repeat step 3) once, remove all liquid from tube without disrupting the beads.
5) Open the cap of non-stick tube, while the tube remains on the magnet, and dry at room temperature for 3 min.
6) Remove the non-stick tube from the magnet, add 46 μL of TE for DNA elution, mix well by pipetting and incubate at room temperature for 5 min.
7) After brief centrifugation, place the non-stick tube on the magnet for 2 min until the liquid clears, transfer all 44 μL of supernatant to a new 0.2 mL PCR tube ready for PCR in next step, or store at -20℃.

### 5. PCR

1)

| Components | Volume |
|---|---|
| DNA | 44 μL |
| PCR Enzyme Mix | 50 μL |
| PCR Primer Mix | 6 μL |
| Total | 100 μL |

2) Place above PCR tube in a Thermal Cycler, and then initiate the reaction as per following conditions:

| Temperature | Time | Cycles |
|---|---|---|
| Heated lid | On | |
| 95℃ | 3 min | |
| 98℃ | 20 sec | 8 |
| 60℃ | 15 sec | |
| 72℃ | 30 sec | |
| 72℃ | 10 min | |
| 4℃ | Hold | |

### 6. Purify PCR Product

1) Place AMPure XP magnetic beads at room temperature 30 min in advance, mix well

by vortexing before use.

2) Add 100 μL of AMPure XP magnetic beads to 100 μL of PCR product, mix well by gently pipetting 10 times, and incubate at room temperature for 5 min.
3) After brief centrifugation, place the non-stick tube on the magnet for 2 min until the liquid clears, remove and discard the supernatant with a pipette.
4) Add 500 μL of fresh 80% ethanol, while the tube remains on the magnet, then, rotate the tubes in the rack by half turns 4 times to wash the beads. Carefully remove and discard the supernatant after 1 min.
5) Repeat step 4) once and try to suck up all liquid from tube bottom.
6) Open the cap of non-stick tube, while the tube remains on the magnet, and dry at room temperature for 3 min.
7) Remove the non-stick tube from the magnet, add 32 μL of TE water for DNA elution, mix well by pipetting and incubate at room temperature for 5 min.
8) After brief centrifugation, place the non-stick tube on the magnet for 2 min until the liquid turning clear, transfer the supernatant to a new non-stick tube. Proceed next step reaction or store at -20℃.

### 7. Homogenization

1) Use double-strand DNA quantification kit such as Qubit® dsDNA HS Assay Kit or Quant-iT™ PicoGreen® dsDNA Assay Kit, and quantify the sample as per the instructions of the quantification kit.
2) It is recommended to mix samples of different Barcodes here.
3) Add mixed sample 300ng (calculated based on its concentration) to a PCR tube, then add NF water to final volume of 48 μL.

### 8. Circularization

1) Denature the homogenized PCR product on a Thermal Cycler at 95℃ for 3 min, then immediately transfer to ice batch.
2) Prepare reaction mixture on ice as per following system:

| Components | Volume |
|---|---|
| Splint Buffer | 11.6 μL |
| Ligation Enzyme | 0.2 μL |
| Total | 11.8 μL |

3) Add 11.8 μL of above reaction mixture to 48 μL of denatured DNA.
4) Place above PCR tube in a Thermal Cycler, and initiate the reaction as per following conditions:

| Temperature | Time |
|---|---|
| Heated lid | on |

| | |
|---|---|
| 37℃ | 30 min |
| 4℃ | Hold |

### 9. Digestion

1) Prepare digestion reaction solution on ice as per following system:

| Components | Volume |
|---|---|
| Digestion Buffer | 1.4 µL |
| Digestion Enzyme | 2.6 µL |
| Total | 4 µL |

2) After the circularization reaction is finished, directly add 4 µL of digestion reaction solution into circularized DNA solution, mix well and briefly centrifuge, then place the PCR tube in a Thermal Cycler, and initiate the reaction as per following conditions:

| Temperature | Time |
|---|---|
| Heated lid | On |
| 37℃ | 30 min |
| 4℃ | Hold |

3) Add 7.5 µL of Digestion Stop Buffer to each reaction, mix well to terminate the reaction.
4) Transfer all the reaction solution to a new non-stick tube, ready for purification.

### 10. Purify Digestion Product

1) Place AMPure XP magnetic beads and place at room temperature for 30 min in advance. Mix well by vortexing before use.
2) Pipette 168 µL AMPure XP magnetic beads to digestion product, mix well by pipetting 10 times, and incubate at room temperature for 10 min.
3) After transient centrifugation, place the non-stick tube on the magnet for 2 min until the liquid clears, remove and discard the supernatant with a pipette:
4) Add 500 µL of fresh 80% ethanol, while the tube remains on the magnet, then, rotate the tubes in the rack by half turns 4 times to wash the beads. Carefully remove and discard the supernatant after 1 min.
5) Repeat step 4) once and try to suck up all liquid from tube bottom.
6) Open the cap of non-stick tube, while the tube remains on the magnet, and dry at room temperature for 3 min.
7) Remove the non-stick tube from the magnet, add 32 µL of TE for DNA elution, mix well by pipetting and incubate at room temperature for 10 min.

8) After brief centrifugation, place the non-stick tube on the magnet for 2 min until the liquid turning clear, transfer the supernatant to a new non-stick tube. Store at -20℃, ready for preparation of DNB.

Ivanova, N.V., Dewaard, J.R. & Hebert, P.D. (2006) An inexpensive, automation-friendly protocol for recovering high-quality DNA. *Molecular ecology notes,* **6,** 998-1002.
