## Supplementary material for "Access COI barcode efficiently using high throughput Single-End 400 bp sequencing": Supplementary File 2.pdf

### HIFI-SE Pipeline User's Guide

Generating COI barcodes using BGISeq-500 Single-End 400 bp sequencing reads (SE400 module)

https://pypi.org/project/HIFI-SE/

Version: 1.0.2; December 10, 2018

Chentao Yang, Guanling Meng
BGI-Shenzhen, Shenzhen 518083, China
China National GeneBank, BGI-Shenzhen, Shenzhen, 518120, China

Copyright (C) 2018 BGI-Research at Shenzhen.

Permission is granted to make and distribute verbatim copies of this manual provided the copyright notice and this permission notice are retained on all copies.

HIFI-SE is licensed and freely distributed under the GNU General Public License version 3 (GPLv3). For a copy of the License, see http://www.gnu.org/licenses/.

### INTRODUCTION

#### Design goal of HIFI-SE

HIFI-SE is designed to handle Single-End 400 bp long sequencing reads generated form latest BGISeq-500 SE400 module, including raw reads filtering, assigning reads to samples, and assembling to COI barcodes. Compared to traditional Sanger COI barcode method, HIFI-SE aims to be significantly more accurate and more able to retrieve COI barcodes from specimens in a cost-effective way, because of the strength of its multiplexing PCR with tagged primers strategy and long sequencing capability.

#### What's still missing in HIFI-SE

Even though this is a stable public release that we consider suitable for most demands, at the same time, HIFI-SE remains a work in progress. We think the codebase is a suitable foundation for development of a number of significantly improved approaches to classically important problems in COI assembly. Some of the more important "holes" to fill for us are:

**Denoising sequencing reads.** Denoising also called noise reduction, is the process of removing noise from a signal. For NGS data, especially amplicons, denoising generally refers to a computational method for removing sequence errors from amplicon reads or, equivalently, identifying the correct biological sequences in the reads. In 16S study, Usearch has provided some powerful tools to denoise PCR reads, such as unoise2 and unoise3, but that was designed for Illumina reads. It is not ideal to transplant UNOISE algorithm in our study. there are still a lot of work to do.

**Multi-process of assembly.** HIFI-SE just provides a single thread mode during filtering, assigning and assembly process expect clustering via Vsearch, because it still achieves an acceptable performance to deal ca. 10G raw reads. Anyway, I will develop a multi-thread version to improve performance further.

**Data visualization.** A statistical result will be presented by a histogram or pie figure in future.

#### How to learn more about HIFI-barcodes study

Actually, HIFI-barcode project is a series study of using high-throughput sequencing platforms (HTS) to establish COI barcode reference. In previous study, we have successfully built the pipeline supplied on Illumina Hiseq platforms. You can get details from previous publication (Liu *et al.* 2017), protocol (dx.doi.org/10.17504/protocols.io.k9icz4e), and software (https://github.com/comery/HIFI-barcode-hiseq; https://github.com/comery/HIFI-barcode-pacbio) or other information in Researchgate (https://www.researchgate.net/project/HIFI-barcode).

### INSTALLATION

#### System requirement and dependencies

**Operating system:** HIFI-SE is designed to run on most platforms, including UNIX, Linux and MacOS/X. Microsoft Windows. We have tested on Linux and on MacOS/X, because these are the machines we develop on. HIFI-SE is written in python language, and a version 3.5 or higher required.

**Dependencies**:

- biopython version 1.5 or higher (required). Please check https://biopython.org/ and https://pypi.org/project/biopython/#description for more details on installation of biopython.
- Another python package - *bold_identification* is also required for getting complete function of HIFI-SE. See https://pypi.org/project/bold-identification/
- HIFI-SE supposed you have installed the VSEARCH on your device, and its path in your $PATH. See https://github.com/torognes/vsearch

#### Installation

Download vsearch and install, to download the **source distribution** from a release and build the executable and the documentation, use the following commands:

```
wget https://github.com/torognes/vsearch/archive/v2.9.1.tar.gz
tar xzf v2.9.1.tar.gz
cd vsearch-2.9.1
./autogen.sh
./configure    # You may customize the installation directory using the --
prefix=DIR option to configure
make
make install    # as root or sudo make install
```

Sometimes, installation from source distribution is not as easy as expected, you can visit google forum to find helpful information about common questions.

Or these commands if you are using a Mac:

```
> wget
https://github.com/torognes/vsearch/releases/download/v2.9.1/vsearch-
2.9.1-macos-x86_64.tar.gz
tar xzf vsearch-2.9.1-macos-x86_64.tar.gz
```

Or if you are using Windows, download and extract (unzip) the contents of this file:

```
https://github.com/torognes/vsearch/releases/download/v2.9.1/vsearch-2.9.1-win-x86_64.zip
```

Linux and Mac: You will now have the binary distribution in a folder called vsearch-2.9.1-linux-x86_64 or vsearch-2.9.1-macos-x86_64 in which you will find three subfolders bin, man and doc. We recommend making a copy or a symbolic link to the vsearch binary bin/vsearch in a folder included in your $PATH, and a copy or a symbolic link to the vsearch man page man/vsearch.1 in a folder included in your $MANPATH. The PDF version of the manual is available in doc/vsearch_manual.pdf.

Now install HIFI-SE, here we provide two ways to install it, installation by pip (python package installation management tool) or copying the source code from github and installing biopython and bold_identification by yourself.

- From PyPi

Installation by pip is recommended because it will solve package dependencies automatically, including biopython and bold_identification packages. Make sure you know what pip is and have you installed pip. Install HIFI-SE for your system using the following commands **(NOTE: pip is link from pip3)**:

```
> pip install HIFI-SE
```

- From Github

I only deploy my latest version on github, so you can clone this repository to your local computer. However, it would not solve package dependencies, thus you need to install biopython and bold_identification before using HIFI-SE software.

```
> git clone https://github.com/comery/HIFI-barcode-SE400.git
> pip install biopython
> pip install bold_identification
```

If you have installed successfully, HIFI-SE.py will be in the directory where your python3 install in, then type HIFI-SE.py in command to test.
Screen shows:

```
usage: HIFI-SE [-h] [-v]
all,filter,assign,assembly,bold_identification ...

Description

    An automatic pipeline for HIFI-SE400 project, including filtering
    raw reads, assigning reads to samples, assembly HIFI barcodes
    (COI sequences).
```

```
Version

    v1.0.2 2018-12-10  Add "-trim" function in filter;
     accept mistmatched in tag or primer sequence,
    when demultiplexing; accept uneven reads to
    assembly; add "-ds" to drop short reads before
    assembly.
    1.0.1 2018-12-2  Add "polish" function
    1.0.0 2018-11-22 formated as PEP8 style
    0.0.1 2018-11-3

Author
    yangchentao at genomics.cn, BGI.
    mengguanliang at genomics.cn, BGI.

positional arguments:
  all,filter,assign,assembly,polish,bold_identification
    all                run filter, assign and assembly
    filter              filter raw reads
    assign              assign reads to samples
    assembly            do assembly from input fastq
                       reads, output HIFI barcodes.
    polish              polish COI barcode assemblies,
                       output confident barcodes.
    bold_identification
                       do taxa identification
                       on BOLD system,

optional arguments:
  -h, --help          show this help message and exit
  -v, --version        show program's version number and exit
```

Add an alias for HIFI-SE.py, type in command:

```
> alias HIFI-SE="HIFI-SE.py"
```

### RELEASE NOTES

## V1.0.2

HIFI-SE v1.0.1 2018/12/2. Changers form previous version:

- add "-trim" function in filter;
- accept mismatch in tag or primer sequence when demultiplexing;

- accept uneven reads to assembly;
- add "-ds" to drop short reads before assembly.

## V1.0.1

HIFI-SE v1.0.1 2018/12/2. Changers form previous version:

- Add “polish” function
- Fixed several small bugs.

## v1.0.0

HIFI-SE v1.0.0 2018/11/22. Changers form previous version:

- Formatted python code writing style as PEP8.
- Fixed several small bugs.

## V0.0.3

HIFI-SE v0.03 2018/11/15. Changes from previous version:

- Modify the description of some arguments for better understanding.

## V0.0.1

HIFI-SE v0.0.1 2018/11/03 beat version, establish the framework and archive almost complete functions.

### TUTORIAL

Here’s a tutorial walk-through of some small projects with HIFI-SE. This should suffice to get you started on work of your own, and you can (at least temporarily) skip the rest of the Guide, such as all the nitty-gritty details of available command line options.

#### The programs in HIFI-SE

**Core commands**

| | |
|---|---|
| all | run filter, assign and assembly |
| filter | filter raw reads |
| assign | assign reads to samples |
| assembly | do assembly from input fastq, reads, output HIFI barcodes. |
| polish | polish COI barcode assemblies, output confident barcodes. |

**Other utility**

| | |
|---|---|
| bold_identification | do taxa identification on BOLD system |

#### Files used in tutorial

All related files could be downloaded from here (https://github.com/comery/HIFI-barcode-SE400/) in `tutorial/` directory. The important files for tutorial are:

– `raw.fastq.gz`, raw output fastq file generated from BGISeq-500 SE400 module.
– `indexed_primer.list`, tagged primer list

#### Run in all

HIFI-SE support an easy way to run entire pipeline by using "all" command. It requires a raw fastq file as input to generate clean reads, then as well as a tagged primer list file to assign reads to samples, finally to assemble reads to COI barcode, respectively. The adapter in raw fastq file must be remove before getting the boll running.

```
> HIFI-SE all -raw raw.fastq.gz -primer indexed_primer.list -index 5 -outpre test
-threads 2
```

If Vserach is not in your environment path, you also can specify Vsearch path using "-vsearch" argument. Like:

```
> HIFI-SE all -raw raw.fastq.gz -primer indexed_primer.list -index 5 -outpre
test   -vsearch /home/your/path/vsearch -threads 2
```

#### Run by steps

- **Step 1, filter raw reads**
  using "-e", expected error threshold to filter, see also chapter 6.

```
> HIFI-SE filter -raw raw.fastq.gz -e 10 -n 3 -outpre test
```

using "-q", Phred quality (known by Q for NGS sequencing data).

```
> HIFI-SE filter -raw raw.fastq.gz -q 20 10 -n 3 -outpre test
```

"-q 20 10" means discarding read which contains more than 10% of quality score < 20 bases.

- **Step 2, assign reads to samples**

```
> HIFI-SE assign -fq test_filter_highQual.fastq -outpre test -index 5 -primer indexed_primer.list
```

- **Step 3, assembly reads to COI barcode**

```
> HIFI-SE assembly -outpre test -index 5 -list test_assign.list -mode 1 -vsearch /home/your/path/vsearch -threads 2
```

**Polish raw assembly results**

```
> HIFI-SE polish -i test_assembly.fasta -index 5
```

**Do taxa identification on BOLD system**

```
> HIFI-SE bold_identification -i test_assembly.fa -o test_assembly.fa.bold.tab -d COX1
```

#### Output files

```
Filter step:
- test_filter_lowqual.fastq
    low quality reads filtered out.
- test_filter_highqual.fastq
    high quality reads kept for assignment
- test_filter_log.txt
    log file of filtering gives a brief report: how many clean reads, how many reads filtered out.
Assign step:
- test_assign.list:
    clean reads path of each sample, one line for a sample. Format like:
    assigned/For001.fastq  assigned/Rev001.fastq
    assigned/For002.fastq  assigned/Rev002.fastq
    …
    assigned/For096.fastq  assigned/Rev096.fastq
- test_assign_assign_err.fasta
```

All reads failed to assign to samples.

- assigned/

  directory contains all assigned reads.

- test.assign.log

**assembly step:**

- test_assembly.fasta

  A standard fasta file format, ID is formatted as follow:

  e.g.

  >004_1-1;1_4;size=2.5;overPos=83;oid=96.39%;len=717

  - 004 is sample id;
  - 1-1 means the consensus sequences of first cluster of 5′ end and first cluster of 3′ end;
  - 1_4 means abundance of consensus form 5′ and 3′ end;
  - size=2.5 means on average abundance of both ends;
  - overPos=83 means overlap length of assembly is 83;
  - oid=96.39% means similarity of overlapped region;
  - len=717 means total length of assembly.

- test_assembly.log

  The head 10 lines starting with "#" show the arguments of user's settings:

  ```
  ## assigned reads list file = test_assign.list
  ## Using all reads to make consensus, no limitation
  ## input read length = 400
  ## vsearch path = vsearch
  ## check codon translation = no
  ## consensus mode = 1
  ## clustering identity = 0.98
  ## overlaping identity = 0.95
  ## min overlap = 80
  ## max overlap = 90
  ```

  The following lines contain the clusters information by number order, which contains input read number and consensus sequence.

- test_assembly.depth

  Each line presents base depth of each site of assembly, e.g.

  004_1-1    1    A:1 T:0 C:0 G:0

  If any two base's depth is same, it will be reported as a heterozygosis base.

When all steps finished, except above files, all failed samples would be listed in "test_rerun.list", then you can run another round with "-rc" argument.

**polish step:**

- test_assembly.fasta.confident

  All COI barcodes have been polished by following conditions:

```
The coverage of 5′ or 3′ end > 5
The length of COI barcode > 650bp
The amino acid translation is good
```

### ARGUMENTS RECOMMENDATION

#### Choosing the maximum expected error threshold

**Phred scores**

Next-generation reads specify a Phred score for each base, also known as a Quality or Q score. The Q score is an integer, typically in the range 2 to 40 (there are several coding systems, such as Sanger, Illumina 1.3+, Illumina 1.5+, Illumina 1.8+, 1.9). Q indicates the probability that the base call is incorrect (P). For example, Q=30 means that the error probability is 0.1%, so the machine is reporting that the base is more likely to be right than wrong.

$$P = 10^{-Q/10}$$

$$Q = -10 \log_{10}(P)$$

**Expected number of errors**

Assume that the Phred scores are good, in other words assume that the machine makes good estimates of the error probabilities. Now we can measure the overall quality of a read by calculating the expected number of errors in the read. The expected value of a quantity under a probability distribution is a technical term in statistics, so let me explain what that means in this case.

Take the error probabilities according to the Q scores in a given read, and imagine a very large number of reads containing errors that occur with those probabilities. For example, if the first base in the read has Q=20, then 1% of the reads in our collection will have errors in that position. Some of the reads will have no errors, some will have one error, and so on. The Q scores can't tell us exactly how many errors there are in a given read. However, the Q scores can tell us what the average number of errors would be in that large collection, and this is the definition of the expected number of errors. Because we're taking an average, the value is real rather than integer, and can be less than one.

**Calculating the expected number of errors and most probable number of errors**

It turns out that it is easy to calculate the expected number of errors: it is the sum of the error probabilities. The most probable number of errors (E*) is also easy to calculate. First calculate E = expected errors = sum P. Then round down to the nearest integer, and this is the most probable number of errors. In symbols: E* = floor(E). For example, if E < 1, then the most probable number of errors is zero. Proofs of these results are given in (Edgar *et al.* 2011)

$$E = \text{expected errors} = \text{sum}(P)$$

$$E^* = \text{floor}(E)$$

**Choosing the maximum expected error threshold**

Edgar, the author of Usearch, recommend that E_max = 1 is a natural choice for Illumina reads, because the most probable number of errors of a filtered read is zero. That make scene because Illumina Q scores are pretty good (much better than current SE400 module). A maximum expected error threshold of 1 will discard many reads that is out our expectation. So I recommend to set a relax value for E_max, even 10 as I used.

#### Clustering or not for assembly?

The experimental protocol in metabarcoding and DNA barcoding studies includes amplification by PCR followed by sequencing, which introduces errors in several ways. Amplification introduces substitution and gap errors (*point errors*) due to incorrect base pairing and polymerase slippage respectively (Turnbaugh *et al.* 2010). PCR chimeras form when an incomplete amplicon primes extension into a different biological template (Haas *et al.* 2011). Sequencing also introduces point errors due to substitutions (incorrect base calls) and gaps (omitted or spurious base calls). Contaminants from reagents and other sources can introduce spurious species (Edgar 2013). Spurious species can also be introduced when reads are assigned to incorrect samples due to *cross-talk*, also known as *tag switching* or *barcode switching (Carlsen et al. 2012)*. However, right now HIFI-SE doesn't provide an ideal solution, instead, we use a clustering method to remove potential errors. I will address this problem in the future. So I recommend you to use "mode=1" to run assembly module.

#### How to run fast?

Here are some tips to speed the program:

- Set a larger threads value for Vsearch
- Set a narrow range of overlap length from -min to -max. e.g. -min=80; -max=85, (while you know, actually overlap length is 83 by and large.)
- Set -seqs_lim to control how many clean reads used to assembly COI barcodes.

### SOME OTHER TOPICS

#### How do I cite HIFI-SE?

Currently this work has not been published, but I will update this part after publication.

#### How do I report a bug?

Before we can see what needs fixing, we almost always need to reproduce a bug on one of our machines. This means we want to have a small, reproducible test case that shows us the failure you're seeing. So if you're reporting a bug, please send us:

- A brief description of what went wrong.
- The command line(s) that reproduce the problem.
- Copies of any files we need to run those command lines.
- Information about what kind of hardware you're on, what operating system, what version of associated packages you used, with what configuration arguments.

Depending on how glaring the bug is, we may not need all this information, but any work you can put into giving us a clean reproducible test case doesn't hurt and often helps. In addition, we're not a company with dedicated support staff – we're a small lab of busy researchers like you. Somebody here is going to drop what they're doing to try to help you out. Try to save us some time, and we're more likely to stay in our usual good mood.

##### Can I use another software to cluster read?

Definitely yes! But before I tell you how to modify a version to meet your demand, I would like to point out that Usearch is not an open source software, but Vsearch is. So, this is the reason why I use Vsearch instead of Usearch.

Let me show you how to modify it.

And first, you should know how and where use cluster software in HIFI, yes, that is in assembly step and just called in mode 1 (clustering reads and make consensus sequence from clusters). In HIFI-SE latest version 1.0.2, from line (L) 958 to line 1067, it defined the function "mode_vsearch", you can change L978-L988 to adapt for Usearch or other software. And before that, remember to modify L552 which is looking for usearch in your $PATH. Importantly, you must set "-uc" to generate a tab-delimited file which contains clusters information, because HIFI-SE will read this file to extract sequence in downstream.

#### MANUAL PAGE

##### filter

usage: HIFI-SE filter [-h] -outpre <STR> -raw <STR> [-e <INT>] [-q <INT> <INT>] [-n <INT>]

| Arguments | type | Descriptions |
|---|---|---|
| -outpre | <str> | Prefix for output files |
| -raw | <str> | Input raw Single-End fastq file, and only adapters should be removed; supposed on Phred33 score system (BGISEQ-500). |
| -e | <int> | Expected error threshold, default=10 see more: http://drive5.com/usearch/manual/exp_errs.html |
| -q | <int> <int> | Filter by base quality; for example: '20 5' means dropping read which contains more than 5 percent of quality score < 20 bases. -e is conflict with -q **(this is important!)** |
| -trim | - | whether to trim 5' end of read, it adapts to -e mode or -q mode |
| -n | <str> | Remove reads containing [INT] Ns, default=1. |

#### assign

usage: HIFI-SE assign [-h] -outpre <STR> -index INT -fq <STR> -primer <STR> [-outdir <STR>]

| Arguments | type | Descriptions |
|---|---|---|
| -outpre | <str> | Prefix for output files |
| -index | <int> | The length of tag sequence in the ends of primers |
| -fq | <str> | Cleaned fastq file |
| -primer | <str> | Taged-primer list, on following format:<br>Rev001 AAGCTAAACTTCAGGGTGACCAAAAAATCA<br>For001 AAGCGGTCAACAAATCATAAAGATATTGG<br>...<br>**this format is necessary!** |
| -outdir | <str> | Output directory for assignment, default="assigned" |
| -tmis | <int> | mismatch number in tag when demultiplexing, default=0 |
| -pmis | <int> | mismatch number in primer when demultiplexing, default=1 |

#### assembly

usage: HIFI-SE assembly [-h] -outpre <STR> -index INT -list FILE
[-vsearch <STR>] [-threads <INT>] [-cid FLOAT]
[-min INT] [-max INT] [-oid FLOAT] [-tp INT] [-ab INT]
[-seqs_lim INT] [-len INT] [-mode INT] [-rc] [-cc]
[-codon INT] [-frame INT]

| Arguments | type | Descriptions |
|---|---|---|
| -outpre | <str> | Prefix for output files |
| -index | <str> | The length of tag sequence in the ends of primers |
| -list | <str> | Input file, fastq file list. [required] |
| -vsearch | <str> | Vsearch path (only needed if vsearch is not in $PATH) |
| -theads | <int> | Threads for vsearch, default=2 |
| -cid | <float> | Identity for clustering, default=0.98 |
| -min | <int> | Minimun length of overlap, default=80, **it can not be more than 83** |
| -max | <int> | Maximum length of overlap, default=90, **it can not be less than 83** |
| -oid | <float> | Minimun similarity of overlap region, default=0.95 |
| -tp | <int> | How many clusters will be used in assembly, recommendation=2 |
| -ab | <int> | Keep clusters to assembly if its abundance >=INT, **-tp is conflict with -ab (this is important!)** |
| -seqs_lim | <int> | Reads number limitation. by default, no limitation for input reads |
| -len | <int> | Standard read length, default=400 |
| -ds | - | drop short reads away before assembly |
| -mode | [1,2] | 1 or 2; modle 1 is to cluster and keep most [-tp] abundance clusters, or clusters abundance more than [-ab], and then make a consensus sequence for each cluster. modle 2 is directly to make only one consensus sequence without clustering. default=1 |

| | | |
|---|---|---|
| -rc | - | Whether to check amino acid translation for reads, default not |
| -codon | <int> | Codon usage table used to check translation, default=5 |
| -frame | [0,1,2] | Start codon shift for amino acid translation, default=1* |

#### Polish

usage: HIFI-SE polish [-h] -i STR [-cc] [-cov INT] [-l INT] -index INT
[-codon INT] [-frame INT]
HIFI-SE polish: error: the following arguments are required: -i, -index

| Arguments | type | Descriptions |
|---|---|---|
| -i | <str> | COI barcode assemblies |
| -cc | | whether to check final COI contig's mino acid translation, default yes |
| -cov | <int> | minimun coverage of 5' or 3' end allowed, default=5 |
| -l | <int> | minimun length of COI barcode allowed, default=650 |
| -index | <int> | the length of tag sequence in the ends of primers |
| -codon | <int> | codon usage table used to checktranslation, default=5 |
| -frame | [0,1,2] | start codon shift for amino acidtranslation, default=1 |

* frame refers to GFF frame, 0 means codon translation start from the first site, 1 means translation starts from the second site, whereas 2 means from third site. For LCO1490 and HCO2198 primer set, it is 1 in most case.
